# The importance of orangutans in small fragments for maintaining metapopulation dynamics

**DOI:** 10.1101/2020.05.17.100842

**Authors:** Marc Ancrenaz, Felicity Oram, Nardiyono, Muhammad Silmi, Marcie E. M. Jopony, Maria Voigt, Dave J.I. Seaman, Julie Sherman, Isabelle Lackman, Carl Traeholt, Serge Wich, Matthew J. Struebig, Truly Santika, Erik Meijaard

## Abstract

Orangutans (*Pongo* spp.) occur at low densities and therefore large areas are necessary to sustain viable metapopulations, defined here as sets of conspecific units of individuals linked by dispersal. Historically, orangutans lived in large contiguous areas of intact rainforest, but are now increasingly found in agricultural and other landscapes modified by people. Here we collate evidence of orangutans utilizing isolated forest fragments (< 500 ha) within multiple-use landscapes dominated by oil palm monoculture across Borneo. Orangutan signs (i.e. nests) were evident in 76 fragments surveyed by helicopter, and in 50 of 70 additional fragments surveyed on the ground; on average 63 ha in size. This includes presence of adult resident females with dependent young confirmed in 40% of the fragments assessed by ground survey. Our study revealed some resident females are raising offspring in isolated forest patches within mature oil palm stands. This not only confirms that some forest patches can sustain orangutans, but indicates migratory males are capable of reaching these fragments scattered throughout the multiple-use landscape. Therefore, orangutans that use or live in even small isolated forest patches are an essential part of the overall metapopulation by maintaining gene flow between, and genetic connectivity within, populations distributed across larger multiple-use landscapes. Orangutan survival is commonly thought to be low in small, isolated forest patches, and the customary management strategy is to remove (translocate) these individuals and release them in larger forests. In some cases, translocations may be necessary, i.e. in case of fire or when the animals are in eminent danger of being killed and have no other refuge. However, the small amount of data available indicates that mortality rates during and after translocations are high, while the impacts of removing animals from spatially dispersed metapopulations are unknown. Therefore, we argue the current policy of routine translocation rather than conserving the species within human-modified landscapes could inadvertently decrease critical metapopulation functionality necessary for long-term viability. It is clear that orangutans need natural forest to survive, but our findings show that fragmented agricultural landscapes can also serve as complementary conservation areas in addition to fully protected areas if they are well designed with ecological connections, and if orangutan killing can be prevented. To achieve this, we call for a paradigm shift from the traditional large single forest model to one that emphasizes metapopulation functionality in the fragmented forest – human use matrix characteristic of the Anthropocene.

Version 2 (20 May 2020): this manuscript is a non-peer reviewed preprint shared via the BiorXiv server while being considered for publication in a peer-reviewed academic journal. Please refer to the permanent digital object identifier (https://doi.org/10.1101/2020.05.17.100842).

Under the Creative Commons license (CC-BY Attribution-Non Commercial-No Derivatives 4.0 International) you are free to share the material as long as the authors are credited, you link to the license, and indicate if any changes have been made. You may not share the work in any way that suggests the licensor endorses you or your use. You cannot change the work in any way or use it commercially.

## 1 Introduction

In international wildlife conservation, the prevailing policies and conservation strategies in governmental and non-governmental organizations have favored large, connected “natural” areas, especially in tropical conservation, such as the “Heart of Borneo”, or the “Central Forest Spine” of West Malaysia. These strategies often consider fragments of natural habitat as of little or no value for wildlife conservation (Sodhi et al., 2010). However, the importance of small habitat patches for biodiversity conservation is increasingly recognized (Edwards et al., 2019; Wintle et al., 2019), especially for wide-ranging species such as large mammals or volant species (Beca et al., 2017; Melo et al., 2017).

Prior to the arrival of modern humans in Asia, orangutans presumably depended on primary forest for survival, but researchers have recently documented their resilience to drastic habitat changes (Spehar et al., 2018). Orangutans persist and reproduce in forests logged for timber (Husson et al., 2009; Ancrenaz et al., 2010), in industrial timber plantations (Meijaard et al., 2010; Spehar and Rayadin, 2017) and in agricultural landscapes, for example (Campbell-Smith et al., 2011). They are also found in isolated patches of forest within landscapes that have been extensively transformed by humans, albeit at lower densities than in more extensive natural forests (Ancrenaz and Lackman, 2014; Ancrenaz et al., 2015; Spehar et al., 2018; Seaman et al., 2019). Orangutan survival and population viability within these heavily managed landscapes is likely contingent on hunting and killing being minimized (Marshall et al., 2006; Husson et al., 2009).

Although orangutans are solitary foragers, they live in an abstract community of known and related individuals, where females are resident and males disperse (Arora et al., 2012). Female orangutans are philopatric and strongly tied to their natal area, and the home ranges of maternal kin often overlap considerably (van Noordwijk et al., 2012). They display extreme long-term site fidelity with the core of their home range and are reluctant to move under normal circumstances (Ashbury et al., 2020). Adult flanged males will aggressively defend an area with females in it, especially when females are sexually active (Spillmann et al., 2016).

Currently, orangutans found in small forest patches are generally perceived as “non-viable”, because of insufficient food, the risk of getting killed by people, or because of fires and logging threatening remaining trees (Sherman et al., 2020). Consequently, many wild orangutans observed in such habitat patches are pre-emptively translocated to nearby forests presumed more suitable for their survival. For example, in Indonesian Borneo between 621 and 1,124 wild orangutans were “rescued” from forest fragments in human-modified landscapes between 2007 and 2017 and translocated to other forest areas (Sherman et al., 2020). Very little is known about the survival rate of individuals following translocation, but rare post-translocation monitoring indicates that translocated orangutans struggle to survive (Sherman et al., 2020). In addition, females released into new forest areas will likely not move far from the point of release (Lokuciejewski, 2018). In areas with existing resident females, competition for food increases, potentially displacing newly arrived females from the release site - or *vice versa* (Marzec et al., 2016).

The impacts on orangutan metapopulation of removing individuals from forest patches are poorly understood, but could intensify the effects of fragmentation and jeopardize long-term viability. Information on the counterfactual – i.e. what would have happened to orangutans had they not been removed from patches – would be useful. In the Kinabatangan area of Malaysian Borneo, for example, orangutans have been recorded regularly for >20 years in most small forest patches available in the oil palm dominated landscape (Ancrenaz et al., 2015, and HUTAN, unpubl. data). Some animals travel through the farmland between forest patches, and some resident females reproduce successfully (Ancrenaz et al., 2015; Spehar et al., 2018). Rather than being completely isolated, these individuals form a larger metapopulation, where conspecific groups of individuals are linked by dispersal (Hastings and Harrison, 1994). The combination of multiple orangutan groups living in forest patches irrespective of their size or protection status is inherently important to the long-term conservation of the species (Voigt et al., 2018).

Here, we build on the experience from the Kinabatangan area and compile evidence of orangutans utilizing forest patches in other human-modified landscapes of Borneo. We refine current understanding and knowledge gaps about the persistence of the species fragmented landscapes, and provide recommendations for conservation management.

## 2 Evidence of orangutans utilizing forest patches

### 2.1 Aerial surveys

Aerial surveys were conducted in 2008, 2012, and 2018 to assess orangutan presence in forest patches across the agricultural landscapes of Kinabatangan, South Sandakan Bay, Segama, Beluran and Sugut. Surveys were conducted from a Bell 206 helicopter at about 60 knots and ca. 100 m above the canopy, following protocols established in Sabah (Ancrenaz et al. (2005). Patches were categorized based on their size (<u>very small</u>: < 5 trees; <u>small</u>: 5 trees to <1 ha; <u>medium</u>: 1-10 ha; large: >10 ha), and orangutan nests and signs of human disturbance recorded (see Ancrenaz, 2015). We limited our aerial investigation to patches <500 ha, which corresponds to the upper limit of a female range in most areas (Singleton et al., 2009). A forest fragment was considered isolated if the closest forest was >500 m away.

The 2008 surveys revealed >500 orangutan nests in 76 small forest patches, <100 ha (including a “patch” with one single tree) that were completely isolated within mature oil palm plantations: Kinabatangan (32 patches; >100 nests); Sandakan Bay (eight patches; >100 nests); Sugut (15 patches; > 150 nests); Beluran (seven patches; >50 nests); Lower Segama (14 patches; >100 nests).

The same route over Kinabatangan in 2012 confirmed 15 of the 32 patches were still present (i.e. 53% of the patches had been cleared) and detected >120 nests in 12 of them (Ancrenaz et al., 2015). In 2018, the repeated survey of Sugut recorded significant deforestation, though all of the seven patches of degraded forest surveyed contained nests (> 60 in total).

### 2.2 Ground and interview surveys

In 2019 and 2020, rapid ground surveys revealed orangutans utilizing forest patches in 11 oil palm estates in Sabah and Kalimantan. Nests were often built on taller trees that offered vantage points, although several were detected in oil palms close to forest. We interviewed plantation workers and managers at these sites about orangutan presence and potential conflicts in their estates, using a previously tested protocol (Meijaard et al., 2011; Ancrenaz et al., 2015). Respondents revealed that in seven estates (four in Sabah, three in Kalimantan) they recognized mothers with their young in and around the same patches (12 occupied by nine females). Two respondents stated the same females (four in total) were regularly observed over 5-10 years. Flanged males and smaller individuals (usually without an infant, which could indicate an unflanged or a young male) were reported walking on the ground in oil palm between forest patches. Crop conflicts were rare, and mostly occurred on young palms <3 years old. Most damage to mature palms occurs within the first two rows of palms located along borders with areas of natural forest. Although it was reported that some damages could impair flower and fruit productivity, estate managers interviewed did not consider damage to mature palms (i.e. above 5 years old) a concern.

To complement the results of our rapid ground surveys in February 2020 we sent a questionnaire to four of the oil palm estates, and to three orangutan researchers working in fragmented forest landscapes. Respondents were asked to categorize forest patches in their landscapes by location and size; as a natural forest corridor(s) to larger forest areas; presence or absence of orangutans, including direct sightings of the individuals and indirect sightings, such as nests or pictures from camera traps. The survey covered nine oil palm landscapes, comprising 81 patches; 11 of those were larger than our size threshold (>500 ha) and are not reported here (although they all supported orangutans).

Camera traps or direct sightings confirmed orangutans in 50 of 70 patches (i.e. 71% of the total) with an average patch size of 57 ha (range: 1-286 ha; SD=72 ha). Signs were detected in 14 of 19 patches <10 ha in size; 18 of 26 patches of 10-50 ha; 9 of 10 of 50-100 ha, and 9 of 15 of 100-500 ha (Figure 1).

**Figure 1:**
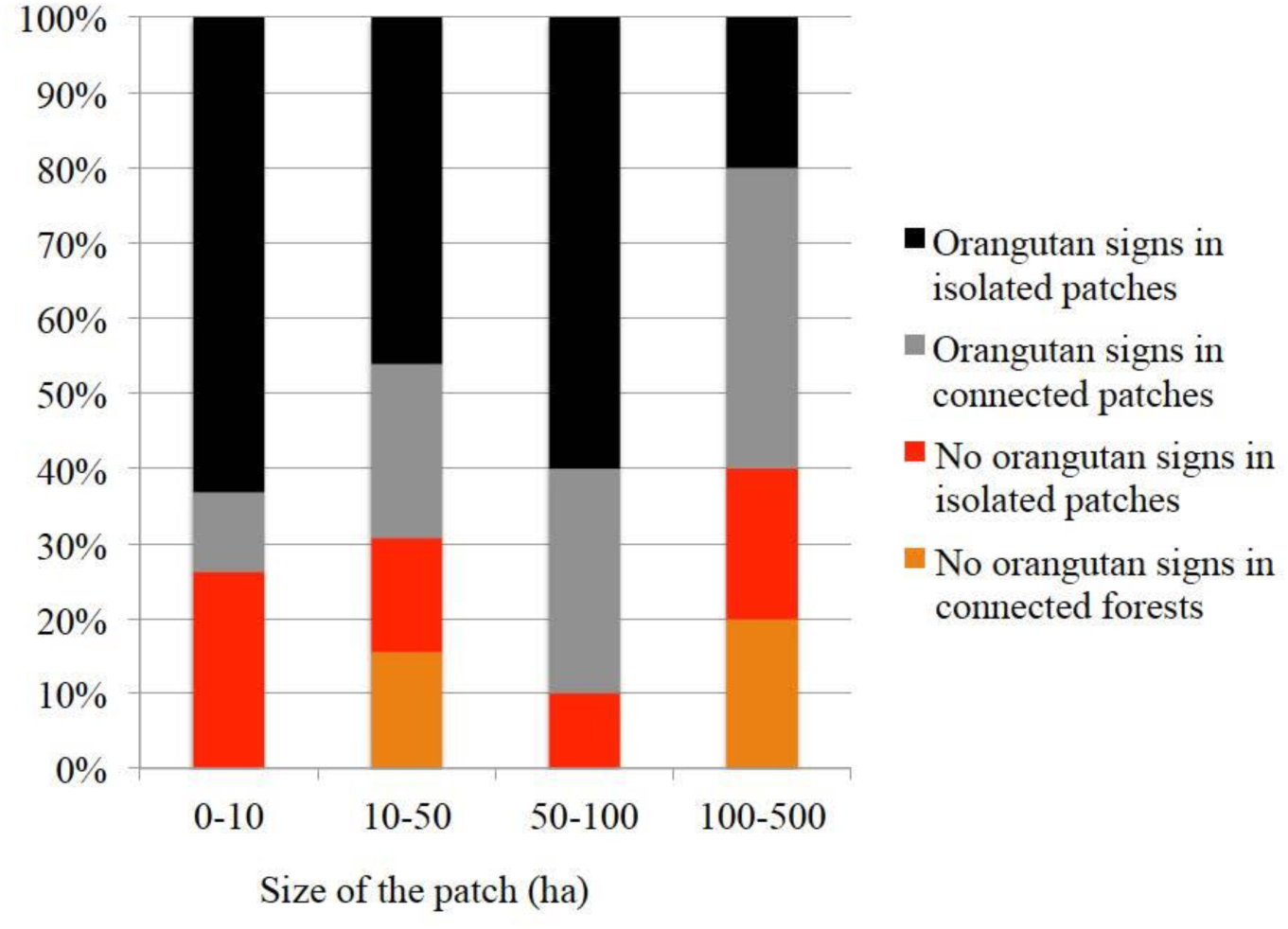
Distribution of orangutan signs in 70 forest patches, according to the size of the fragments (in ha) and their connectivity status with sorrounding forests.

Seventy two percent of patches with orangutan signs were >500 m from the nearest forest and considered “isolated”. Presence of an adult female with young was confirmed in 21 fragments (10-236 ha in size; mean 71; SD= 69), and 12 were completely isolated. In 10 additional patches signs of adult females without young, or unflanged males (these two classes being very difficult to distinguish), were reported. Flanged males were present in four patches, and 15 patches had signs of orangutan nests without any indication of age and sex. Orangutans were absent from 20 patches at the time of surveys (0.5-369 ha in size; mean 81 ha; SD 101 ha), 13 of which were isolated (Figure 1).

Orangutans regularly use forested corridors set aside by estates between forest patches, such as riparian buffers along rivers, or “pathways” designated to enhance connectivity across the landscape. For example, to link two isolated forest patches one estate created a 40 m wide corridor 1.2 km long, comprising trees that had been planted under palms. Orangutan nests were observed in this corridor within four years, confirming use for dispersal. During our site visits, the three large corridors between forests that we assessed had nests, illustrating that corridors are important for resting (nesting) and for dispersing across the landscape.

## 3 Discussion

Our collation of reports from agricultural landscapes demonstrates substantial use of forest fragments by orangutans in farmland. It is increasingly evident that orangutans are a highly flexible and adaptable species that can maintain high population densities in production forests (Ancrenaz et al., 2010; Oram, 2018; Roth et al., 2020), and most wild orangutan populations in Borneo are currently found outside of strictly-protected forests (Santika et al., 2017; Voigt et al., 2018). Orangutans in these landscapes can cope with degradation in habitat structure such as canopy opening (Davies et al., 2017), disperse on the ground when necessary (Ancrenaz et al., 2014), reproduce, and even successfully raise young to maturity (van Noordwijk et al., 2018).

In extensive oil palm plantations, orangutan presence is more prevalent close to forest edges or within patches inhabited by resident individuals, as already documented in Kinabatangan (Ancrenaz et al., 2015). However, both flanged and unflanged males have been recorded to venture several km inside plantations (Ancrenaz et al., 2015). Female orangutans are reluctant to leave their natal area in Bornean forest (Arora et al., 2012; van Noordwijk et al., 2012), and this may also be the case in forest patches: some are not only surviving, but raise offspring in isolated patches many years after planting oil palm. These individuals probably survived the initial phase of forest loss when oil palm estates were established and took refuge in small forest patches retained within the modified landscape. Over the years, they maintained their ranging patterns by visiting and using as many forest patches within their former home-range as possible, even if most of this home-range is currently covered with palms or other agricultural crops (Figure 2). Of course, sufficient food resources are needed for these females to survive and reproduce in these fragmented landscapes. Their chances of long-term survival are likely increased with the number, size and quality of forest fragments. In some places, enrichment planting of key food species, and improving the overall forest connectivity within the agricultural landscape will also be important for increasing survival.

**Figure 2.**
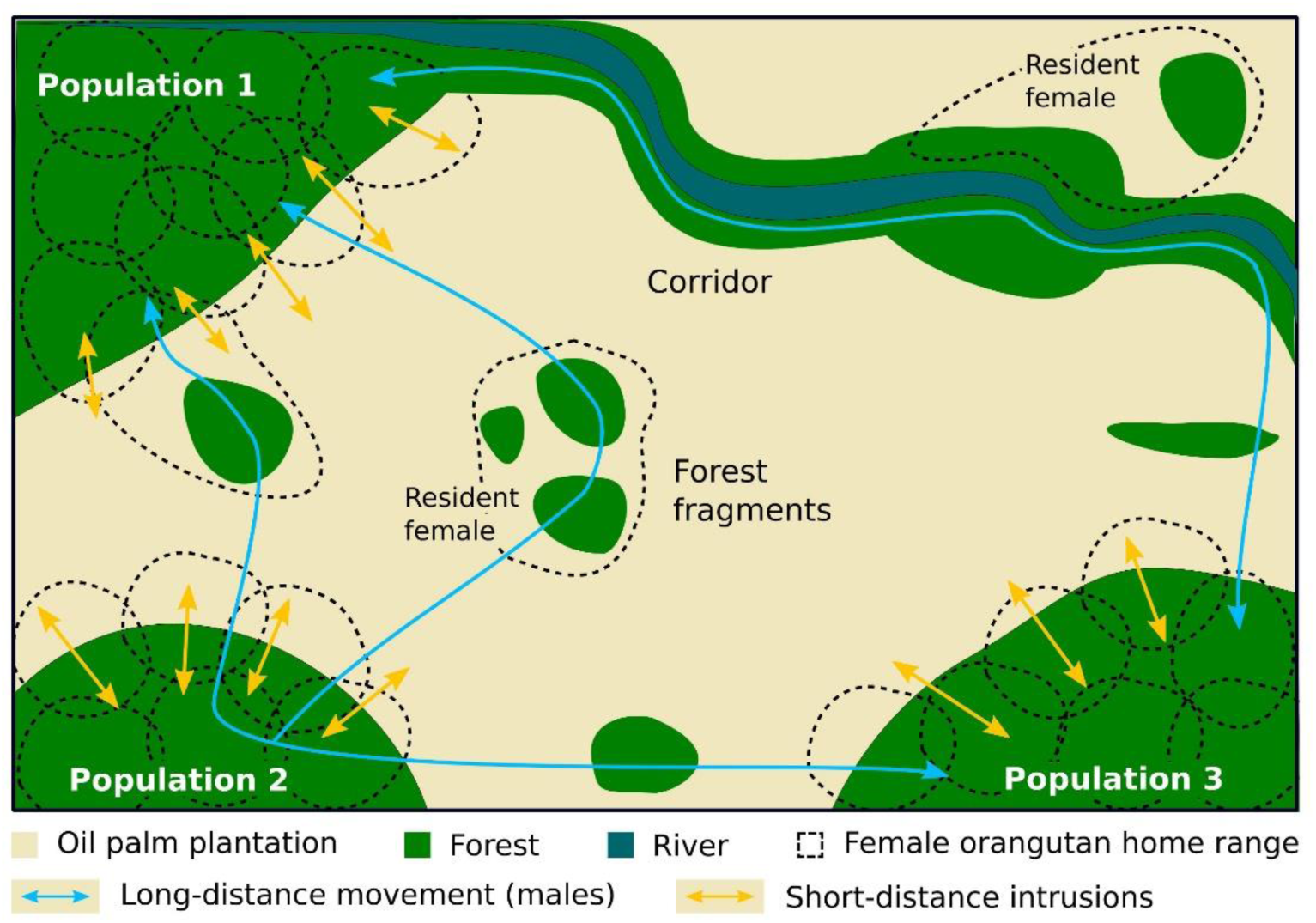
Schematic representation of the metapopulation functioning of orangutans in fragmented landscapes.

The presence of offspring in these isolated patches suggests migrating males are likely using the human-modified landscape to search for receptive females. Indeed, males are the most frequently observed sex in oil palm plantations generally, including numerous mentions of them walking on the ground, along rivers or streams and even on plantation roads. During these travels, orangutans may use any vantage point (such as isolated trees) to navigate within the plantation as suggested by the frequent report of nests built in scattered single trees, or small groups of trees, within plantations far away from any forest. Orangutans also appear to use forest corridors to move across the landscape, as recorded by Seaman et al. (2019) and our surveys in commercially administrated lands. These corridors are often set aside as high conservation value forests to meet sustainability certification criteria, either as riparian buffers or other linkages between forest patches.

Further investigation is warranted before we can consider a primarily agricultural landscape as viable long-term habitat for orangutans. Many questions still need to be addressed, for example: Are orangutan dietary needs met in plantation settings?; What is the fate of young individuals that grow up in small isolated fragments?; How can we get local people and companies to agree living alongside orangutans?; or What are the risks associated with the persistence of fragments as viable habitats (e.g. edge effects, fire, inbreeding, intrusion, etc.) and edge effects? Many questions remain, but it is clear that we need to revisit our thinking about what encompasses a viable orangutan population. The traditional way of thinking of populations as a group of orangutans in a contiguous forest would ultimately result in a disjunct distribution range with isolated populations no longer connected genetically. Long-term maintenance of habitat stepping-stones within larger multifunctional landscapes, on the other hand, could retain connectivity between the larger forest areas and boost the overall chances of survival for the population as a whole. Therefore, conservation efforts for orangutans, and other endangered tropical species, must begin to recognize the critical role habitat fragments may play to help stabilize landscapes for orangutans at the metapopulation level. Government and non-government organizations need to join forces and acknowledge that orangutan survival is best guaranteed through overall management of metapopulations.

Recognizing that orangutan viability hinges on metapopulation integrity also requires a change in perceptions about “rescuing” and translocating individuals. While there are obvious threats to orangutans living in forest fragments (physical harm, killing, or forest destruction), a recent analysis of wild-to-wild translocation in Kalimantan showed that at least 90% of the individuals captured were healthy and several of them had healthy infants as well (Sherman et al., 2020). These animals had managed to survive in habitats perceived as inhospitable by orangutan conservation practitioners. Therefore, we argue that, rather than emptying small forest patches of orangutans as a default operational practice, local authorities and conservation organizations must develop more proactive solutions, where forest patch management fosters measures to allow people and orangutans to co-exist safely. We also argue that removing individuals (especially resident females) could be deleterious to the overall metapopulation functioning of fragmented populations; it may threaten both the orangutan metapopulation functionality at the translocation site, as well as be potentially catastrophic not only to the individual, but the metapopulation at the release site as well. There are at least 10,000 orangutans in multiple use landscapes (Meijaard et al., 2017) and rescuing all is unfeasible, thus requiring *in situ* management given its protected status. It also recognizes that is neither non-zero during wild-to-wild translocations (Wilson and McMahon, 2006), nor following reintroduction (Galdikas, 2018; Sherman et al., 2020).

An additional key problem with translocations is that once the orangutans are removed from a forest patch (or at least those animals that could be captured), the forest patch and its other remaining wildlife are more likely to be converted to human use, because the forest patch has lost what little protection it received from containing orangutans as a legally protected species. Indeed, the presence of a fully protected species in a forest fragment confers to this fragment a status of “High Conservation Value” with a specific set of management measures, including no-deforestation requirements, and this reduces the likelihood of conversion (Carlson et al., 2018). If not protected, the loss of the forest patch would then also mean the loss of all other wildlife that was not rescued as well as loss of ecosystem services provided by the forest (Lucey et al., 2014; Wells et al., 2016).

Of course, there remain circumstances when the health of animals surviving in small fragments is compromised (e.g. food scarcity, habitat destruction (K. Sanchez, pers. com.), and in such cases, translocations will still be needed when the alternative is a dead orangutan. Our paper is not against translocation as a part of the overall conservation toolbox, but we emphasize that this kind of intervention should be the exception rather than the norm. Governments tend to support translocations as it provides a means for them to allow developments to go ahead in unprotected forest areas (Meijaard, 2017), and it is important that the conservation community respects the legal principle that orangutans are protected species, whether they occur inside or outside protected areas. Ongoing discussion with government authorities is needed to ensure that the focus of orangutan conservation strongly remains on *in situ* protection of remaining populations.

## 4 Conclusion

Insights from our current studies require that we reassess the notion of orangutan population viability in human-modified landscapes. Orangutan populations in a contiguous forest area containing fewer than 50 individuals are generally thought to be non-viable (Utami-Atmoko et al., 2019), and as a result are often relocated. However, such a policy overlooks the fact that (1) this is a species that exists at low densities even in ideal conditions, and therefore functions as a metapopulation or set of conspecific groups of individuals linked by dispersal across wide distances; and (2) that in today’s reality on Borneo and Sumatra many forest fragments are not large enough to contain more than 50 animals. The good news, as we demonstrate here, is that metapopulations are still functioning in mixed-use landscapes so gene flow is still possible. In other words, most populations across fragmented landscapes could be viable if we manage to maintain essential habitat fragments and prevent any unnatural deaths or removal from the landscape. Therefore, the conservation unit to be managed should then not be the animals in relatively well-protected larger forest areas, but the metapopulation that is ranging across the entire mixed protected–privately administered landscape as a whole. Eventually, the future of orangutans in the Anthropocene will primarily depend on the attitude of all land users and government that should target a peaceful coexistence between people and orangutans outside and inside protected areas.

## Acknowledgments

In addition to all HUTAN supporters we thank the United States Fish and Wildlife Service Great Ape Conservation Fund for financial support [grant number F17AP01081], the Arcus Foundation, New York, NY [grant number G-PGM-1610-1985], the IUCN Species Survival Commission Primate Specialist Group - Section on Great Apes [project grant number P02472], the Yayasan Sime Darby and the French Alliance for Preservation of Forests. The funders had no involvement in study design, in the collection, analysis and interpretation of data, or in the writing of the paper and the decision to submit the article for publication.

## Author contributions

MA, FO, IL and EM initially conceptualized this study. FO, NA, DS, MJ, MS, Simli, and CT contributed data. MV helped with data analysis and diagram design. MA, EM, FO, JS, CT, MS, MV, and DS helped improve the manuscript.

## Data Availability Statement

The datasets analyzed for this study can be found in the XXXX.

